# Interacting brains revisited: A cross-brain network neuroscience perspective

**DOI:** 10.1101/2021.02.20.432051

**Authors:** C. Gerloff, K. Konrad, D. Bzdok, C. Büsing, V. Reindl

**Affiliations:** Child Neuropsychology Section, Department of Child and Adolescent Psychiatry, Psychosomatics and Psychotherapy, Medical Faculty, RWTH Aachen University, Germany; JARA-Brain Institute II, Molecular Neuroscience and Neuroimaging, RWTH Aachen & Research Centre Juelich, Germany; Chair II of Mathematics, Faculty of Mathematics, Computer Science and Natural Sciences, RWTH Aachen University, Germany; Department of Biomedical Engineering, McConnell Brain Imaging Centre, Montreal Neurological Institute, Faculty of Medicine, McGill University, Montreal, Canada; Mila – Quebec Artificial Intelligence Institute, Montreal, Canada

## Abstract

Elucidating the neural basis of social behavior is a long-standing challenge in neuroscience. Such endeavors are driven by attempts to extend the isolated perspective on the human brain by considering interacting persons’ brain activities, but a theoretical and computational framework for this purpose is still in its infancy. Here, we posit a comprehensive framework based on bipartite graphs for interbrain networks and address whether they provide meaningful insights into the neural underpinnings of social interactions. First, we show that the nodal density of such graphs exhibits nonrandom properties. While the current analyses mostly rely on global metrics, we encode the regions’ roles via matrix decomposition to obtain an interpretable network representation yielding both global and local insights. With Bayesian modeling, we reveal how synchrony patterns seeded in specific brain regions contribute to global effects. Beyond inferential inquiries, we demonstrate that graph representations can be used to predict individual social characteristics, outperforming functional connectivity estimators for this purpose. In the future, this may provide a means of characterizing individual variations in social behavior or identifying biomarkers for social interaction and disorders.

## Introduction

Network models have provided new insights into the neural basis of social behavior. Traditionally, neuroscience studies have examined functional connectivity within the brain of a single subject using standardized tasks. These studies have yielded mounting evidence that the integrative processes and dynamic interplay among different brain regions and systems underpin many of our cognitive processes (*1*). While these studies have focused on single brains, each person is embedded in a larger network of social partners and continuously affects and is affected by others (*2, 3*). Thus, to reach a deeper comprehension of the neural underpinnings of social interactions, it may not be sufficient to study individual brains in isolation (*3*). Instead, the continuous reciprocal exchange between interacting brains should be the target of investigation.

Studies that record the brain activities of two or more persons concurrently, a mode of investigation referred to as hyperscanning, indicate that the brain activities of interacting persons become synchronized, i.e., show statistical dependencies (interpersonal neural synchrony, INS) (*4*). However, many of these studies have analyzed INS only between region pairs of the two participants, and often in homologous brain regions, thereby neglecting the fact that each brain is organized and operates as a complex system [e.g., (*5–7*)]. Such pairwise comparisons of region pairs may not be sufficient to fully capture INS, given the large number of dynamic dependencies between brain regions as well as individual variability in functional networks. While the first few, mostly electroencephalography (EEG)-based, hyperscanning studies have adopted a graph analytic approach to study INS [e.g., (*8, 9*)], graph analysis for hyperscanning data is still in its infancy (*10*). Here, we posit a comprehensive analytical framework for inference and prediction based on bipartite interbrain graphs. Specifically, we extend current analyses by i) considering both global and nodal topological properties, ii) comparing the properties of interbrain graphs to those of random graphs, and iii) exploring whether graph representations can be used to infer relationships and predict social characteristics.

A bipartite graph can be constructed from two sets of nodes representing each person’s brain regions and edges encoding their statistical dependencies. Importantly, such a bipartite graph enables full preservation of spatial precision in two nonoverlapping region-of-interest (ROI) sets, which sets it apart from traditional seed-based connectivity mapping, in which the seed ROI must be averaged, causing its spatial information to be lost. From such a bipartite graph, both global and nodal properties can be derived. Previous graph-based hyperscanning studies have often focused on global metrics [e.g., (*8, 9*)]. A global metric, such as the global efficiency, is a single metric (scalar) that aggregates a specific graph topological property across the entire graph (*11*). Hence, global metrics cannot provide insights into the local differences in a graph’s topology and may thereby overlook smaller and/or more localized effects. Thus, understanding how connectivity varies across nodes is an important step in network analysis (*12*). One of the most fundamental nodal metrics, to which most other network metrics are ultimately linked, is the nodal degree, i.e., the number of edges per node, or the nodal density, i.e., the degree normalized with respect to the number of possible edges (*13*).

### Local topology of real-world networks

Nodal metrics not only provide deeper insights into network organization but were also used in early network neuroscience studies to assess the general properties and feasibility of graph formulations [e.g., (*14*)]. In an intrabrain network, as in many other large, real-world networks, not every node is equally important to the network’s topology. While most nodes have very few connections (a low degree), some nodes have many connections (a high degree), facilitating integration across the network (*12, 15*). Consequently, the degree distribution shows heavy-tailed characteristics and differs from the degree distribution of a random graph with uniform edge probabilities, such as an Erdős-Rényi graph (*16*). However, these fundamental topological properties have so far not been examined in hyperscanning studies, although they may provide first evidence for the validity of a network-based perspective on INS. Thus, in this study, we investigate whether interbrain networks show similar heavy-tailed properties, indicating the presence of seed regions, and importantly, whether their nodal density distributions differ from those of random graphs.

### Bayesian inference based on interbrain network representations

Nodal metrics offer an in-depth view of a graph’s topology. However, they come at the cost of several methodological challenges: while global metrics provide scalar values as summary measures, nodal metrics take the form of vectors of measures. Thus, nodal metrics can be analyzed 1) by formulating one model for each node, leading to problems associated with multiple comparisons, or 2) by formulating a single multivariate model that considers the dependencies among nodes. While the latter approach is superior, it becomes analytically unfeasible with an increasing number of nodes and the resulting increase in model complexity. Hence, an approach that can yield localized, interpretable insights into interbrain networks while avoiding multiple comparison issues is currently lacking. To address this problem, we use an unsupervised machine learning technique, the non-negative matrix factorization [NMF; (*17*)], to decompose an interbrain network into several collections of nodes. This allows us to encode the nodal topologies in a low-dimensional vector while preserving the contributions of the individual nodes to each component. The coefficients of these components are then analyzed within a single multivariate Bayesian hierarchical model (BHM), thereby integrating all nodal effects into a single model and avoiding multiple comparisons. Furthermore, Bayesian analysis goes beyond the point estimates of a frequentist approach by allowing a probability statement to be derived for each parameter at both the individual and population levels [see also (*18*)] and allowing heavy-tailed response distributions to be specified.

### Interbrain network based prediction

The Bayesian model explicitly follows the explanatory goal of inference for building and testing neuroscientific claims [see also (*19*)]. While many previous hyperscanning studies have used inferential modeling, an important long-term goal may be to predict behavioral responses, clinical phenotypes or treatment outcomes. In addition, predictive modeling provides a straightforward way to quantify the performance of our NMF-based approach by comparing its ability to predict known properties of the data to the corresponding capabilities of other approaches. This enables us to address whether interbrain networks provide an informative representation that goes beyond functional connectivity estimators.

Prediction based on vectorized connectivity estimators [see ‘connectivity fingerprinting’ (*20*)] captures only pairwise dependencies while neglecting the multiple dependencies among regions of the same network. In contrast, predictive models for classifying graph-structured data represent a new direction in neuroscience (*21*). The underling computational problem of such graph comparison is well known in graph theory; formally, under the premise to determine whether two graphs are topologically equivalent, termed graph isomorphism problem (*22*). Going beyond first attempts, such as the pairwise Weisfeiler-Lehman test, graph representation learning (GRL) is gaining enormous traction (*23, 24*). The aim of GRL is to learn low-dimensional continuous vector representations for graph-structured data, called embeddings (*25, 26*). These methods involve embedding nodes or graphs that are similar with respect to a certain association into a graph space and then transforming them into Euclidean space (*26*). Since our NMF-based approach preserves the topological properties of a graph, it can be understood as a ‘structural embedding’ of a graph. However, since our approach is based on topological properties, other network embeddings based on higher-order proximities could potentially yield superior network representations. Hence, in a rigorous testing regime, we compare the performance of our network representation with state-of-the-art network embeddings [Graph2Vec (*27*), GL2Vec (*28*), GeoScattering (*29*), LDP (*30*)] and predictions based on functional connectivity estimators.

We demonstrate the proposed analytical framework and evaluate its utility using a functional near-infrared spectroscopy (fNIRS) hyperscanning dataset with a well-established hyperscanning design [for more details, see (*7*)]. fNIRS measures the hemodynamic response in cortical brain regions, usually with a higher spatial resolution than EEG, and is based on changes in the concentrations of both oxy- and deoxyhemoglobin (HbO and HbR). The latter property is an advantage of fNIRS because it allows us to conduct analyses based on the HbO signals, as is most commonly done [e.g., (*5, 7*)], while using the HbR signals as a hold-out dataset for the prediction task to validate the findings.

In the current study, *N* = 34 female children (*M* = 14.26 years, *SD* = 2.21 years) performed both a cooperative and a competitive computer task (with two task blocks each) with their mother and with an adult stranger, while their prefrontal brain activities were recorded concurrently. The two task conditions were compared to a baseline condition in which both participants watched a relaxing video together. Thus, after exclusion of noisy datasets, a total of *n* = 330 graphs based on HbO signals and *n* = 330 graphs based on HbR signals were constructed. Previous research based on the same experimental task enables a plausibility check of the results: based on previous findings, stronger INS is expected for mother-child dyads than for stranger-child dyads (*7, 31*) and for the two task conditions compared to the baseline condition in the dorsolateral and frontopolar brain regions (*31*).

## Results

### 2.1 Inference based on global graph properties

To study interbrain networks and their global properties, we first formulated a complete bipartite graph for each dyad by calculating the pairwise statistical dependencies between neural signals from adult and child regions (see workflow Fig. 1). One well-known issue in graph analysis is that complete graphs are sensitive to errors in the derived edges, leading to spurious connections and high false positive rates (*32*). Hence, complete graphs are often reduced via proportional thresholding; however, this has been shown to preserve some spurious connections, particularly in datasets with a low overall functional connectivity (*32*). Thus, instead of using a proportional threshold, we reduced the complete graphs, deriving thresholds from ‘shuffled’ adult-child pairs. By adapting subsampling-based reduction (*33*) and deconfounding (*34*), shuffled adult-child pairs were constructed by permuting adults and children who participated in the same experimental condition but independently of each other. This enabled us to control for confounding variables during graph reduction by defining an exchangeability assumption for blockwise permutation, i.e., under which conditions the data could be permuted. Specifically, the graphs were reduced by setting the channel combination and condition as fixed and treating the subject IDs as exchangeable. For each edge, an individual edge-weight distribution was derived, and its 95% quantile was set as the threshold. Setting the channel combination as fixed allowed us to control for systematic differences between channels, such as differences in the signal-to-noise ratio related to the distance between the brain and skull or the amount of hair. Furthermore, setting the experimental condition as fixed allowed us to control for systematic differences related not to social interaction per se but rather, for instance, to differences in activation patterns [see also (*31*)].

**Fig. 1.**
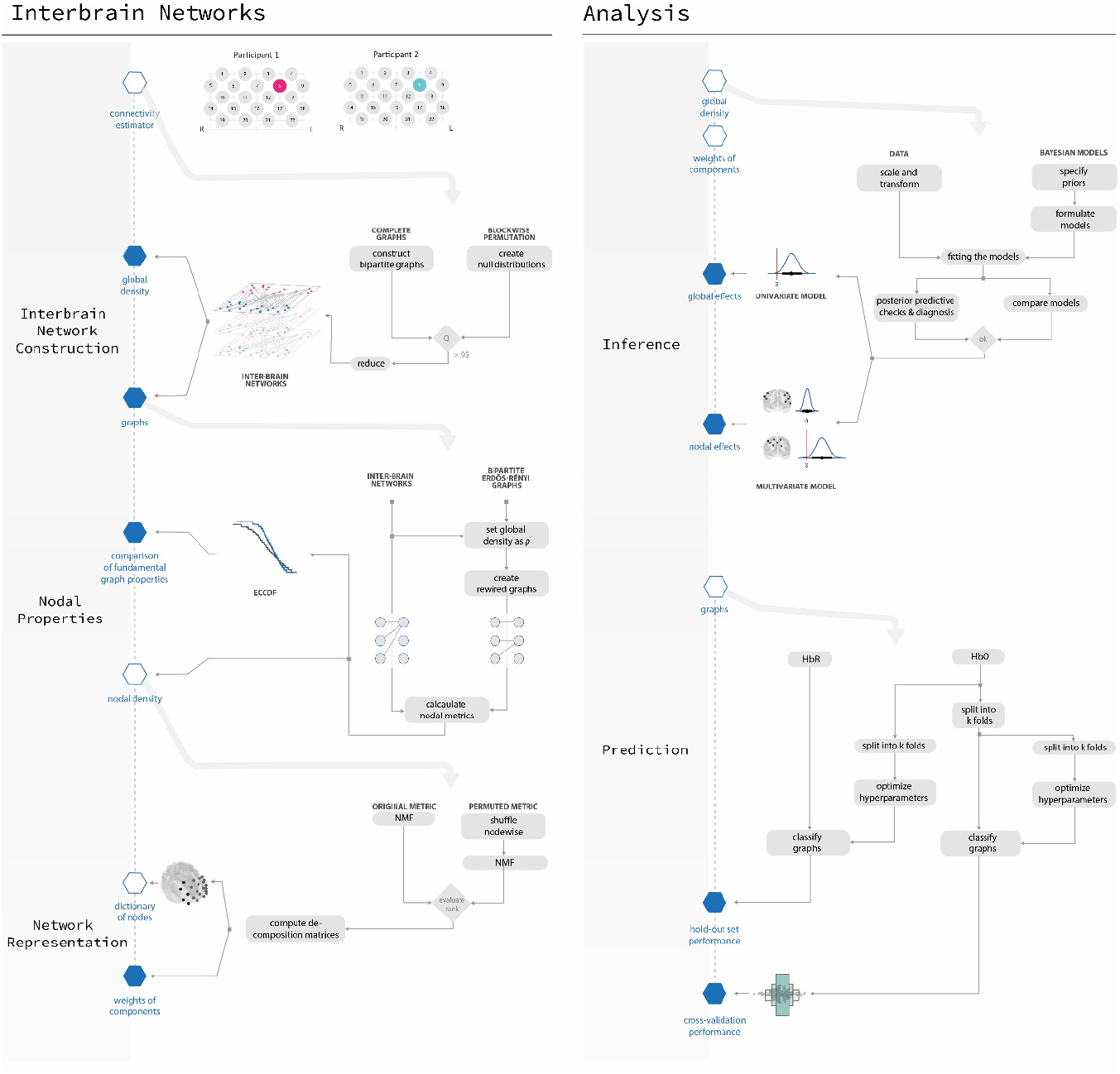
Framework and workflow for inference and prediction based on interbrain networks. The proposed framework governs the construction (*left*) and analysis (*right*) of interbrain networks. For each dyad and condition, a bipartite graph was constructed from the functional connectivity estimators. The complete bipartite graphs for each dyad were reduced via blockwise permutation using distributions obtained from noninteracting dyads (‘shuffled pairs’) to reduce the influence of spurious edges and to stratify for confounds. From each complete bipartite graph, a bipartite Erdős-Rényi graph was constructed while preserving the graph’s global density, and the nodal properties of the original and constructed graphs were compared. Next, the nodal densities of the interbrain networks were decomposed via NMF to obtain an interpretable, lower-rank representation. Analysis of the interbrain networks in both the inference and prediction regimes was demonstrated (*right*). For inference, BHM analysis was introduced, which extends the point estimates of classical frequentist approaches by quantifying the uncertainty of effects using a probability term. First, the global densities of the individual graphs were analyzed to quantify partner and task effects (global effects). Second, the coefficient matrix of our NMF-based network representation was used to formulate a multivariate BHM to yield a more detailed and region-specific quantification of the partner and task effects (nodal effects). Finally, we demonstrated prediction based on interbrain networks using nested-stratified cross-validation of HbO signals and an HbR hold-out set. The performance of our NMF-based network representation was compared with that of state-of-the-art network embeddings and with that of prediction based directly on the functional connectivity estimators.

The global density was then quantified as the average number of edges connected to a node, after correction for noisy fNIRS channels, which were excluded from the analyses. The global densities of the resulting graphs were analyzed within a single BHM with competition (0 = baseline, 1 = competition), cooperation (0 = baseline, 1 = cooperation), and partner (0 = stranger, 1 = mother) as well as the two-way interactions between task (competition/cooperation) and partner as predictors. The results are presented in Fig. 2 and table S1 (Model 1) via the posterior mean (μ), the two-sided 90% credible interval (CI), and the percentage of samples above zero of the marginal posterior distribution. We found evidence for a competitive task effect and, to a weaker degree, evidence for a cooperative task effect, with higher density in the two task conditions compared to the baseline condition (baseline vs. competition: μ = 0.13, CI = [0.02, 0.25], 96.93% > 0; baseline vs. cooperation: μ = 0.08, CI = [-0.02, 0.18], 91.38% > 0). Further, strong evidence was found for a partner effect, with higher density for mother-child dyads compared to stranger-child dyads (μ = 0.13, CI = [0.04, 0.23], 99.3% > 0).

**Fig. 2.**
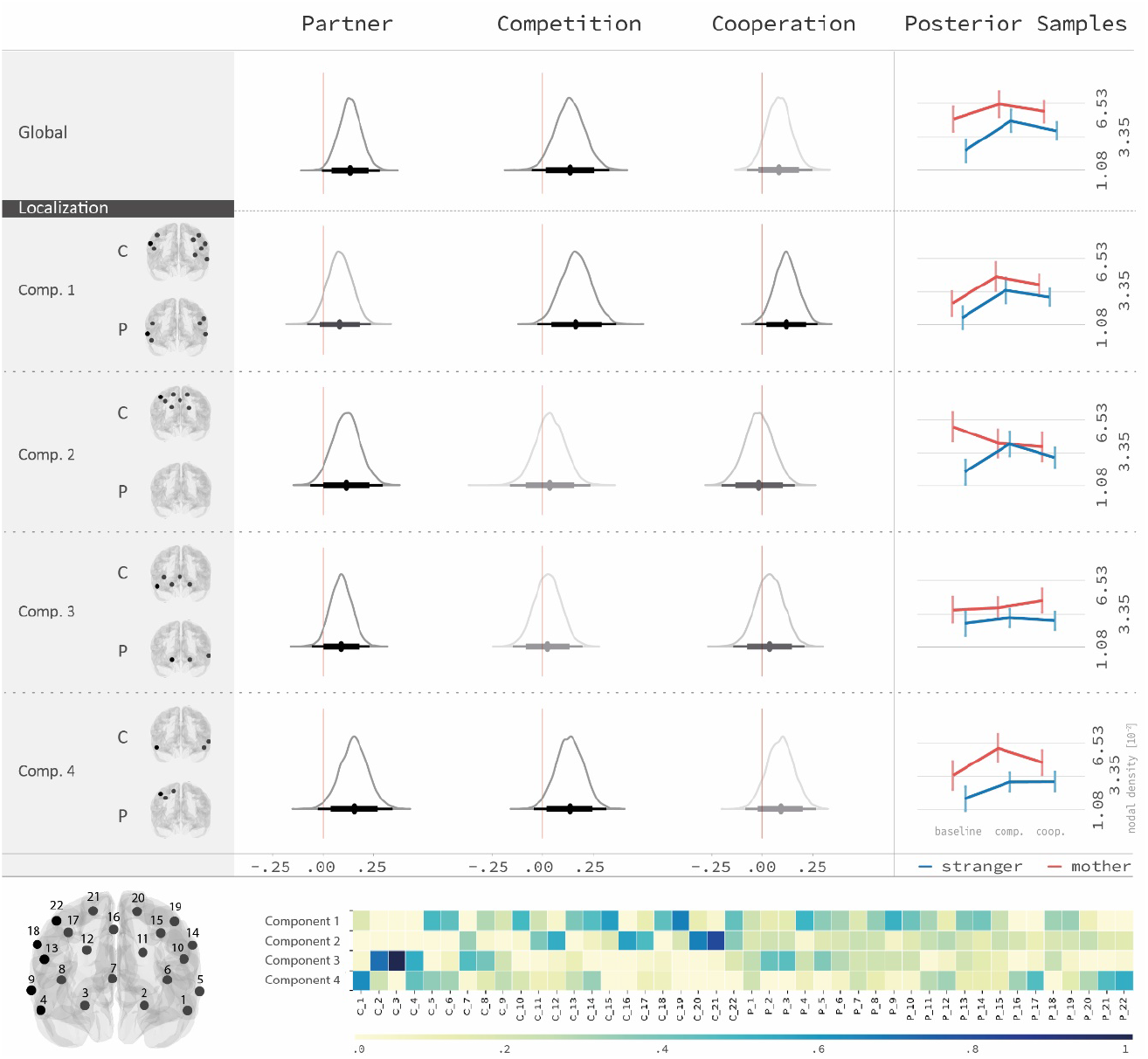
Global effects of interbrain networks can be decomposed into nodal effects in specific seed regions. This figure shows the global and nodal density effects of stranger-child interaction vs. mother-child interaction, baseline vs. competition and baseline vs. cooperation. The nodal densities were encoded by the coefficients of the four NMF components. *Left*: The marginal posterior distributions of the population effects are plotted along with their means, 90% CIs (thick black lines) and 99% CIs (thin gray lines). The width of the CI represents the uncertainty associated with the estimated parameters. Parameters for which the 90% CI does not include zero (red line) were interpreted as evidence for an effect. The global results showed a strong partner effect, with increased synchrony for mother-child dyads compared to stranger-child dyads, as well as a competition effect and some evidence for a cooperation effect, with higher synchrony for the two task conditions compared to the baseline condition. The nodal results confirmed these global results but provided increased topological detail. Specifically, the partner effect was rather widespread (in components 3 and 4 as well as for the baseline condition, in component 2), while the competition and cooperation effects were mostly localized to left and right lateralized prefrontal brain regions (components 1 and 4). *Right*: Five catplots show how the posterior global and nodal density values vary between mother-child and stranger-child dyads in the baseline, cooperation and competition conditions. *Bottom*: A heatmap visualizes the basis matrix resulting from NMF, showing the contribution of each fNIRS channel of the child (C) and the adult partner (P) (x-axis) to the corresponding component (y-axis). The fNIRS channels of the child and the adult partner that contribute most to each of the components in terms of their nodal densities, with weights above the 80th percentile (min = 0, max = 1), are visualized on the brain models.

### 2.2 Nodal properties of interbrain networks

In the previous section, we described the global effects of interbrain networks. Building on this, nodal graph metrics may yield further insights into the specific brain regions that facilitate such effects. Since the nodal degree distribution in a complex system has been shown to follow a more heavy-tailed distribution than that in a random graph with uniform edge probabilities (*13, 15*), we compared the nodal density distributions of the interbrain networks to those of random bipartite Erdős-Rényi graphs. To rule out the possibility that differences these distributions could arise due to differences in global density, we adopted the bipartite Erdős-Reny graphs so that their rewiring regime preserved the global density of the interbrain networks. In the following, the graphs are compared across all experimental conditions, while separate analyses for the partner and task effects can be found in the supplement, text S1, table S2, fig. S1.

Descriptive results showed that the interbrain networks have both a higher proportion of low-density nodes (density ≤ 0.01; 83.42% vs. 81.05%) and a higher proportion of high-density nodes (density ≥ 0.5; 3.34% vs. 1.49%) than Erdős-Rényi graphs with the same global density. These higher proportions of both low-density and high-density nodes indicate that the interbrain networks are more heavy-tailed (Fig. 3). These differences between the nodal distributions of the interbrain networks and Erdős-Rényi graphs were confirmed by a significant result from the two-sided Kolmogorov-Smirnov test (0.117, p < 0.001). To summarize, our results indicate that the distributions of interbrain graphs were more heavy-tailed than distributions of bipartite Erdös-Renyi graphs with the same global properties.

**Fig. 3.**
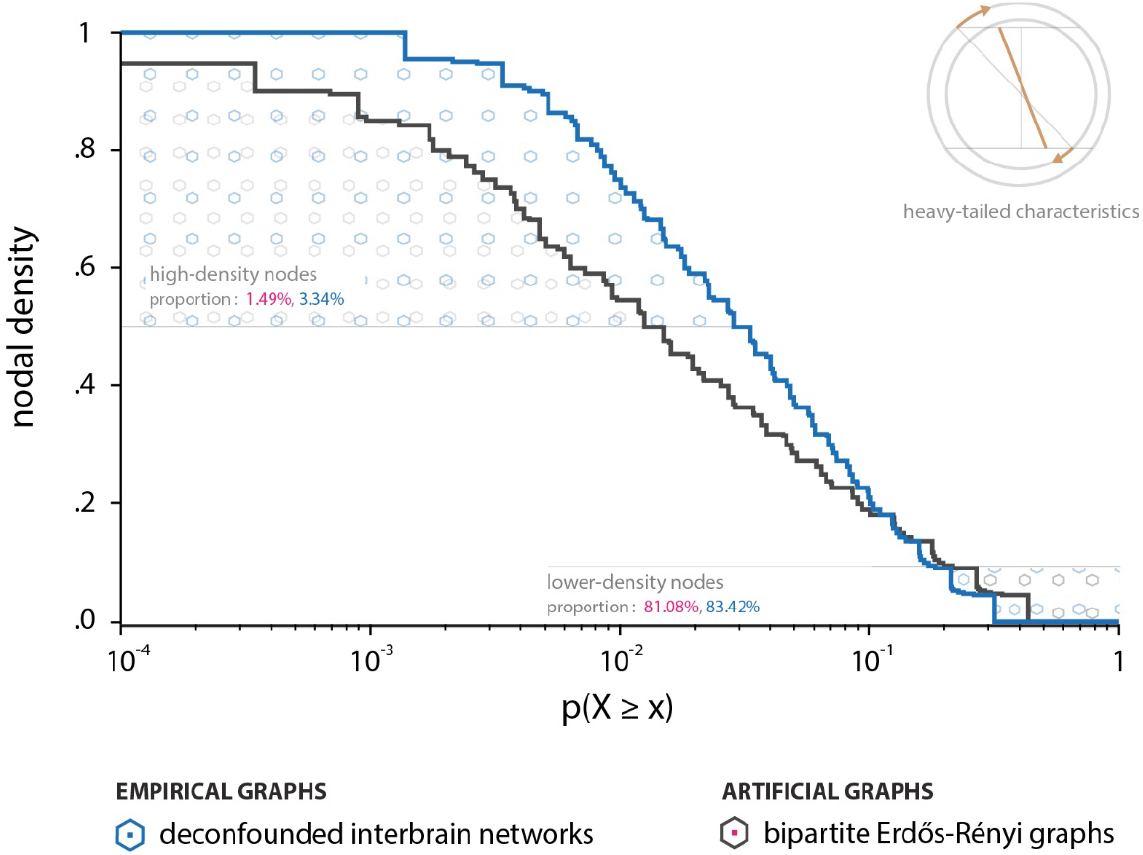
Interbrain networks show nonrandom characteristics typical of many real-world networks. The depicted empirical complementary cumulative distribution functions (ECCDFs) show the tail behavior of the nodal density distributions of interbrain networks (*blue*) and bipartite Erdős-Rényi graphs (*black*). The ECCDF describes the probability with which a node reaches a specific density. As illustrated, the higher steepness of the interbrain ECCDF curve compared to the bipartite Erdős-Rényi ECCDF curve implies a heavier-tailed nature of the distribution, meaning larger proportions of high- and low-density nodes. Such heavy-tailed characteristics are known to be a fundamental property of many real-world networks.

### 2.3 Inference based on nodal graph properties

To analyze the nodal densities while avoiding multiple comparison issues, we derived subsets of nodes showing similar behaviors, reducing the dimensionality of the nodal metrics of each deconfounded graph to four components via NMF. The NMF procedure yields two matrices. The basis matrix can be understood as a dictionary in which to look up the contributions of each node to each of the components that are constant across dyads and conditions (Fig. 2), whereas the coefficient matrix encodes the nodal densities of each dyad in the respective condition. To analyze the nodal densities, a multivariate BHM was again calculated to examine the effects of cooperation, competition and partner as well as the two-way task x partner interactions on the four NMF components.

The nodal density results confirmed the global results, showing evidence for partner, competition and cooperation effects (Fig. 2 and table S1, Model 2). Specifically, NMF component 1 showed strong evidence for both task effects, with increased INS for cooperation (μ = 0.12, CI = [0.02, 0.22], 97.7% > 0) and competition (μ = 0.16, CI = [0.04, 0.29], 98.56% > 0), as well as weaker evidence for a partner effect (μ = 0.08, CI = [-0.02, 0.18], 91.31% > 0), with increased INS for mother-child dyads compared to stranger-child dyads. Component 2 showed evidence for task x partner interactions (competition x partner: μ = −0.27, CI = [-0.49, −0.05], 2.07% > 0, cooperation x partner: μ = −0.21, CI = [-0.44, 0.02], 6.32% > 0), indicating that mother-child dyads exhibited a higher density than stranger-child dyads in the baseline condition. Component 3 did not show any strong task effects but showed evidence for a partner effect, with higher synchrony of mother-child dyads (μ = 0.09, CI = [0.00, 0.18], 95.58% > 0). Component 4 showed a strong partner effect (μ = 0.15, CI = [0.04, 0.27], 98.44% > 0), a competition and a cooperation effect (baseline vs. competition: μ = 0.13, CI = [0.02, 0.25], 97.48% > 0; baseline vs. cooperation: μ = 0.09, CI = [-0.02, 0.20], 90.92% > 0). The brain regions that contribute most to each of the components can be found in Fig. 2. The obtained NMF solution was validated by showing that i) the brain regions that contribute to the NMF components show the same or similar effects as the NMF components themselves (text S2) and ii) the procedure is not permutation biased, an essential requirement in the graph space, meaning that the results are independent of the ordering of the nodes (text S3).

### 2.4 Exploring predictive applications

As described above, the NMF components were first used in an inference task. Next, we explored whether our network representation can also be applied for a predictive goal. To this end, we trained and tested classifiers to discriminate between graphs according to the task conditions (baseline/task), whereby cooperation and competition are summarized to one category (‘task’) and compared to baseline, and the partner conditions (stranger/mother). Four state-of-the-art network embeddings (Graph2Vec, GL2Vec, GeoScattering, LDP) and our NMF-based network representation served as features in the predictive models. Common machine-learning classifiers, namely, an L2-regularized logistic regression classifier [ridge classification (*35*)], gaussian process (*36*), and a support vector machine with linear kernel (*37*)], were applied to compare the performance of the embeddings. Additionally, prediction was performed directly on the functional connectivity estimators. To establish a rigorous evaluation setting, the models were benchmarked via out-of-sample classification performance in the following two regimes:

i. A repeated, stratified, nested k-fold cross-validation procedure was applied based on graphs derived from the HbO signals.
ii. In a true hold-out regime, the classifiers were trained on all HbO graphs and tested on graphs derived from the HbR signals.

First, we assessed whether the predictive models were able to discriminate task conditions (baseline/task) in the two evaluation settings (i, ii). For i), GL2Vec and our network representation showed results that were better than chance across all classifiers (Fig. 4). However, neither method was able to uphold its performance on the hold-out set. GeoScattering was superior in regime ii) but showed performance below chance on i) for the Gaussian process classifier. Prediction based on the functional connectivity estimators did not reach the level of chance for any classifier in both regimes i) and ii).

**Fig. 4.**
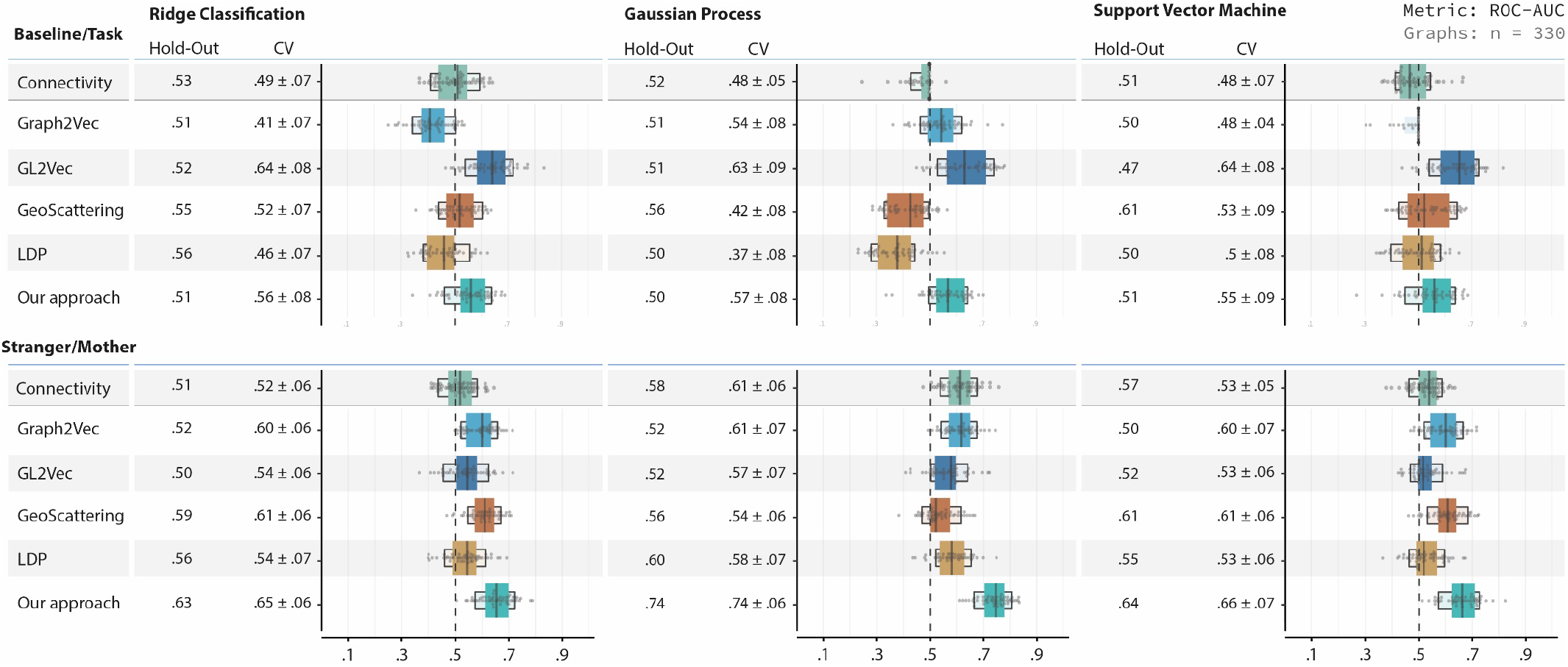
Network representations of interbrain networks might improve the classification of interaction partners and tasks. Classification performance based on network representations and functional connectivity was compared using three widely applied classifiers. For the two classification tasks, baseline/task (top) and stranger/mother (bottom), out-of-sample performance was evaluated via the ROC-AUC. The evaluation regimes consisted of nested cross-validation (*CV; k_out_* = 5, *k_inner_* = 3) and an additional hold-out set (HbR). The CV was stratified to preserve the class distribution across folds and was repeated (n = 10) to avoid the bias that may arise from the splitting of the folds. The ‘letter-value plots’ and the table display the means, standard deviations, and quantiles of the CV performance. To quantify each embedding’ s ability to generalize across datasets, models were trained on the HbO data and used to predict task and partner conditions based on the HbR data. The obtained ROC-AUC results for the CV and hold-out set regimes showed that our proposed NMF-based network representation can successfully discriminate between partners (stranger/mother). GL2Vec showed better performance for task prediction (baseline/task), although task prediction generally appeared more challenging, particularly on the hold-out data. In addition to the network embeddings, the same analysis was performed directly on the functional connectivity estimators. The results showed that prediction based on network representations generally outperformed prediction based on functional connectivity, supporting the validity of using interbrain networks for the analysis of INS.

Second, we evaluated whether the graphs could be classified based on the partner condition (stranger/mother). Consistent with the partner effect evidenced by the BHM analysis (see sections 2.1 and 2.3), most embeddings showed results better than chance. Generally, the performance was higher for partner classification than for task classification across all classifiers, with the exception of GL2Vec. Our NMF-based approach outperformed all network embeddings as well as the functional connectivity estimators across all classifiers in both evaluation settings (i, ii). Again, GeoScattering reached competitive results in regime ii). Both GeoScattering and our approach showed comparable performance between i) and ii). GL2Vec showed relatively consistent performance better than chance in the cross-validation regime (i) but exhibited poor generalization across datasets (ii) (fig. S4B). A closer look at the recall of Graph2Vec, revealed that it tended to assign a majority of graphs to the same class (fig. S4B). Detailed results, including additional performance metrics, classifiers, and parameters, can be found in the fig. S4 and table S3.

## Discussion

Extending the isolated view of a single brain by examining the statistical dependencies between interacting brains, may shed new light on the neurobiological underpinnings of social interactions. While network analysis is an established, state-of-the-art tool for investigating brain connectivity, hyperscanning studies often analyze only the pairwise relationships between the brain regions of one participant and the brain regions of another. Moving beyond this pairwise analysis, we here present the first comprehensive analytical framework for inference and prediction based on bipartite interbrain networks. Specifically, we demonstrated that the nodal densities in interbrain network are characterized by skewed, heavy-tailed distributions. To gain local insights, we showed how global effects can be decomposed via NMF into several individual effects within subsets of nodes. These nodal effects were analyzed within a single BHM to avoid multiple comparison issues while considering skewed response functions. Beyond inferential inquiries, we also introduced prediction based on individual interbrain networks. These first results suggest that network embeddings, and our NMF-based approach in particular, provide informative representations of INS whose predictive power exceeds that of the pairwise statistical dependencies measured on the basis of connectivity estimators.

### 3.1 Bipartite graphs as a representation of interbrain synchrony

In hyperscanning studies, INS is often calculated region-by-region between homologous brain regions [e.g., (*5–7*)]. However, this approach implicitly assumes that synchrony arises primarily in homologous brain regions, thereby neglecting the fact that each brain is organized and operates as a complex network. Along these lines, mentalizing in more neutral social interactions may elicit activity in a broader network than the activity elicited in response to static, isolated stimuli. This may be the case either because this broader network may be characteristic of how the brain processes real-world social stimuli or because multiple processes are simultaneously engaged (*38*). In the cooperative and competitive tasks, these processes may include prediction and adaption processes, emotional process, social comparison processes but also processing of stimuli and shared attention to the task [see also (*7, 31*)]. Thus, information processing in social interactions is complex and likely goes beyond activity occurring in a single, circumscribed brain region, but most likely engages multiple, interacting brain networks in both subjects. Examining synchrony only in a pairwise, region-by-region fashion may fall short of capturing this complexity. This general critique of pairwise analyses is grounded in system theory, as such analyses ignore the multiple dependencies among the entities of a system (*23*).

In addition to neuroscientific arguments for a graph-based approach, there are also several methodological advantages. First, research shows that interindividual differences exist in both the brain structures and networks in which cognitive, emotional, and social processes are substantiated [e.g., (*39, 40*)]. These differences may be even more pronounced when studying dyads of different age groups, such as adults and infants/children, considering the ongoing maturation of brain structures and development of functions in the latter group [see also (*41*)]. Such inter-individual differences may obscure effects in a pairwise (region-by-region) analysis. On top of such anatomical differences, uncertainties in channel localization may arise due to the spatial resolution of the technique, as both fNIRS and EEG are inferior in this respect to functional magnetic resonance imaging (fMRI).

To summarize, we propose to move away from a pairwise analysis and towards a graph-based analytical approach when measuring INS between interacting persons for the reasons described above, namely, i) the engagement of brain networks rather than single brain regions, ii) inter-individual anatomical differences, and iii) uncertainties in localization. Yet, this system perspective comes with several analytical challenges, such as how to identify and deal with spurious edges (see section 2.1) and how to gain local insights into a network (see section 3.3).

### 3.2 Nodal properties of interbrain networks

Investigating the properties of interbrain networks, we have shown that the nodal density distribution is more heavy-tailed than the distribution of random graphs with the same global density. Such skewed distributions can be found on nearly all physiological and anatomical levels of the brain as well as in many other networks - encountered in fields ranging from biology to social sciences to infrastructure (*42, 43*). Thus, these results indicate that interbrain networks have properties similar to those of many complex real-world networks and are significantly different from random graphs. In intrabrain networks, such high-density nodes may act as network hubs, which promote the integration of information and efficient neural communication (*1*). While the origin and function of high-density nodes in interbrain networks are less understood, both intrabrain hubs and interbrain seed regions play an important structural role in their respective network topologies due to their high densities [see also (*44, 45*)]. These findings have important practical implications for future research. First, many classical frequentist tests require a Gaussian distribution and thus are ill suited for analyzing heavy-tailed distributions. Second, this study provides the first evidence that some nodes have unproportionally high degrees, which implies that focusing on global metrics alone is not sufficient to examine how brain activities synchronize during social interactions. In contrast, a nodal analysis not only may yield more topological details but could also reveal more localized effects that would otherwise be obscured when considering global metrics alone. Finally, these findings should be considered a first starting point that raises new questions about the role of high-density regions in interacting yet structurally unconnected brains. For instance, future research may address whether and how high-density nodes in inter- and intrabrain networks are related to each other.

### 3.3 Inference based on interbrain networks

We have successfully demonstrated that interbrain networks can be constructed in the form of bipartite graphs. Regarding the global density, our BHM results show that the density was higher for the two interactive conditions in our study (cooperation/competition) than for the noninteractive baseline condition. In line with our previous findings in male children and adolescents (*31*), this indicates that interbrain networks can encode interaction specific neural processes distinct from those associated with a noninteractive, rest-like state. Moreover, the mother-child dyads showed higher INS than the stranger-child dyads. This was a general effect, observed across the baseline and task conditions, suggesting that the brain responses may be more in tune with each other due to genetic factors, the dyad’s affiliation or shared past experiences. This finding is consistent with several other studies showing increased INS in close relationships when performing joint motor tasks (*6, 7, 31, 46*). For a more detailed discussion of the task and partner effects see (*47*).

Subsequently, nodal properties were compared, which provide more precise, localized information than global graph metrics or alternative subgraph-based approaches (*48*). To reduce the dimensionality, we encoded the nodal properties via NMF. The NMF is applied in various domains and to address a variety of problems, such as the detection of communities or the clustering of neural connections (*49, 50*). One of its key advantages is its ability to extract sparse and meaningful features. While there are nearly infinite possibilities to decompose a matrix or picture, NMF allows to identify latent factors that can be interpreted as ‘parts-of-whole’, such as eyes within a picture of a person (*17*).

Analogously, here, we found that the NMF was able to decompose the global density effect into subsets of nodes showing particular distinct result patterns. Specifically, we found cooperative and competitive task effects in components 1 and 4, located mostly in the left and right lateralized dorsolateral and frontopolar brain regions of adult and child, as well as evidence for a partner effect in components 3 and 4, located mostly in the left, right and medial frontopolar brain regions of both partners as well as the right dorsolateral brain regions of the adult [for further discussion of the brain regions see (*47*)]. Thus, the NMF results confirmed the global results but provided additional topological details. Importantly, we were able to show that our NMF based method is not permutation biased and thus fulfills an essential requirement for graph representation.

### 3.4 Classification performance of network embeddings

Next, we extended the application of the NMF-based method from inference to prediction to answer the question of whether network representations have the capacity to predict experimental conditions on an individual level. Although all network representations achieved a performance above the chance level for predicting the interaction partner (stranger/mother), the NMF-based representation presented here showed the best performance across both a rigorous cross-validation regime and prediction on a second dataset (HbR hold-out set). In contrast to our NMF-based representation, graph embeddings often rely on higher-order proximities, i.e., similarities that can be derived from complex operations on the adjacency matrix, at the cost that they are difficult or impossible to interpret. This makes them suitable for predictive tasks but less so for inference tasks (*51*). In contrast, a few embeddings, such as LDP (*30*) exploit the possibility that nodes may play similar structural roles within the network. LDP is therefore most similar to our approach but considers the structural role of a node by its degree, whereas our NMF-based representation is based on nodal density, which is less biased by excluded channels, and considers more complex, additive patterns. LDP showed an area under the receiver operating characteristic curve (ROC-AUC) above chance level and yielded somewhat competitive results compared to other embeddings but was consistently outperformed by our NMF-based approach. This finding supports the concept of encoding the brain regions’ structural roles but shows that considering more complex, additive patterns and a graph metric that is not biased by excluded channels may be advantageous. Importantly, it was found that both GeoScattering, a state-of-the-art graph embedding based on adjusted wavelet transformation for graphs, and the NMF-based representation could be trained on one dataset and successfully ported to another dataset, indicating reasonable generalization across diverse settings. Such inhomogeneous datasets are likely to arise in neuroscience due to different experimental designs, possible confounders or reduced standardization in naturalistic settings. Consequently, to elucidate the neural basis of social interaction, generalization across datasets will probably be a challenging key factor for predictive applications.

While partner classification showed promising results, task classification was challenging. GL2Vec showed the best performance and reasonable behavior in terms of precision and recall in the cross-validation setting (Fig. 4 and fig. S4A). Interestingly, GL2Vec, which is based on a line graph (*52*), achieved relatively consistent performance above the chance level for cross-validation but showed poor generalization across datasets. A line graph provides an edge-centric representation because it maps the edges of the original graph onto the nodes of the line graph. Recent studies [e.g., (*53*)] suggest that edge-centric approaches may be advantageous in considering individual differences in brain networks. It can be speculated that the interesting properties of line graphs which may better capture individual variances could also potentially make generalization across inhomogeneous datasets more challenging. Overall, the lower performance for task classification may be explained by the class imbalance between the baseline (*n_baseline_* = 64) and task (*n_task_* = 266) conditions. Very high and low precision and recall values, respectively, indicate that especially Graph2Vec but also our NMF-based approach might suffer from class imbalance and, for some instances, classify the majority of graphs only as task or only as baseline (fig. S4A). Although we have made some effort to tackle the imbalance (text S9), low performance may be driven by the fact that not all dyads showed an increased INS during task conditions.

A central finding from our analyses pertains to the general capacities of network representations. Network representations performed at least as well as or better than predictions based directly on the functional connectivity estimators. These results indicate that network representations of INS may carry further information compared to the ‘pure’ functional connectivity estimators. Importantly, in contrast to the other embeddings, our NMF-based approach not only yielded competitive results but also preserved interpretability.

### 3.5 Limitations

Although the current study provides the first evidence that the proposed graph analytical framework can be successfully applied to studies on interbrain connectivity, a number of caveats must be considered. First, intrabrain relationships were not explicitly modeled in the current study, a limitation that could be addressed by other graph formulations. Second, the current reduction procedure enables stratification for confounds but does not ensure a weakly connected graph without isolated nodes, i.e., nodes with no connection to any node. Thus, it cannot be applied to network embeddings that require all nodes to be connected. Third, while our results show that network representations could potentially serve for predictive tasks in hyperscanning, the baseline/task classification task in particular presents a challenge that will require further improvements, which are beyond the scope of this paper. Furthermore, no appropriate benchmark datasets are available for GRP for interbrain networks, and thus, the results presented here should be considered only a first starting point. Nevertheless, the promising performance of some embeddings in addition to our own approach motivates the development of new, more comprehensive benchmark datasets to better evaluate and analyze different methods of GRL.

### 3.6 Conclusions

Network models may contribute to a better understanding of the neural underpinnings of social interaction, possibly providing a more precise analytical operationalization of the concept of INS than pairwise statistical dependencies. Based on bipartite graphs, we have proposed a novel computational approach to capture the dynamic interplay of brain regions that considers the multiple dependencies across the brain regions of interacting persons. The heavy-tailed characteristics of these interbrain networks, differing from those of random graphs, as well as the replication of previous findings regarding partner and task effects and the demonstrated predictive power of interbrain network representations argue for their validity. This framework comes with several advantages: i) it yields both global and nodal insights; ii) through Bayesian modeling, it avoids the problems associated with multiple comparisons, accounts for skewed distributions, and provides uncertainty estimates; and iii) it is applicable for both inference and prediction tasks while providing interpretable insights. Thus, the proposed framework has potentially far-reaching implications, as it enables the exploration of the specific brain regions in a network that seed neural synchrony between interacting persons - an approach that could also be applied for the topological analysis of intrabrain networks. Above and beyond inferential inquiries, our results are the first to demonstrate that network embeddings could provide an informative representation of interpersonal synchrony and may thereby stimulate further research into prediction based on graph-structured data. In the long-run, these models could have important clinical applications, such as predicting treatment outcomes.

## Materials and methods

### 4.1 Empirical data

#### 4.1.1 Participants

Data were collected as part of a larger project, described in (*47*). Participants were recruited via previous studies, postings on the intranet of the University Hospital RWTH Aachen and flyers, and they were screened for severe cardiac, neurological or psychiatric conditions prior to study inclusion. The final sample consisted of 34 children (all female) who participated in the study with their mothers (mother-child dyads) and a previously unacquainted female adult (stranger-child dyads). A total of 29 strangers were included, some of which participated twice (*n* = 1) and some three times (*n* = 2), thus, comprising 34 mother-child and 34 stranger-child dyads. The average age of the children was 14.26 years (*SD* = 2.21 years, range: 10 – 18 years), that of the mothers was 45.32 years (*SD* = 4.95 years, range: 37 – 56 years), and that of the strangers was 23.07 years (*SD* = 2.09 years, range: 19 – 29 years). The strangers were significantly younger than the mothers (*t* (61) = −22.532, *p* <.001). All adults gave written informed consent for their own study participation and, in the case of the mothers, for the participation of their children. The children gave written informed assent (aged 10 – 17 years) or informed consent (aged 18 years). Ethics approval was obtained from the Ethics Committee of the Medical Faculty, University Hospital RWTH Aachen (EK 151/18).

#### 4.1.2 Data acquisition

fNIRS signals were recorded from both members of each dyad concurrently using a single fNIRS device (ETG-4000, Hitachi Medical Corporation, Japan; sampling rate: 10 Hz). Fifteen probes, eight sources and seven detectors, were arranged in grids consisting of three rows and five probe columns, in which the source-detector separation was fixed at 3 cm and the sources and detectors were arranged in alternating order, resulting in 22 measurement channels (CHs). Probe grids were mounted on modified EEG caps (Easycap GmbH, Germany) and placed symmetrically on the participant’s foreheads, with the middle probe of the lowest probe row placed on the Fpz point in the 10-20 system and the middle probe column aligned along the sagittal reference curve. This probe setup, depicted in Fig. 2, covered parts of Brodmann areas (BAs) 8, 9, 10 and 46. The most likely Montreal Neurological Institute (MNI) coordinates of the probes and channels as well as their underlying brain regions were estimated via the virtual registration method (*54, 55*), using the Talairach Daemon (*56*) [for the most likely MNI coordinates of the fNIRS channels see: (*57*)]. For more details on the data acquisition, see (*58*).

#### 4.1.3 Tasks and procedure

The participants in each dyad were seated next to each other in front of a single computer screen. A towel was placed over their hands to reduce their ability to view each other’s movements. Furthermore, the participants rested their chins on chin rests to reduce head movements and improve the fNIRS data quality. The experiment consisted of three experimental conditions: a noninteractive baseline, a cooperative task and a competitive task. Each child completed the three conditions either first with the mother (*N* = 17, 50%) and then, after a short break, with the stranger or the other way around (*N* = 17, 50%). The three conditions were presented in the same order for both dyads of which each child was a member, always starting with the baseline condition, followed either by the cooperative (*N* = 16, 47%) or the competitive (*N* = 18, 53%) task.

##### Baseline

A three-minute excerpt from a relaxing aquatic video (Coral Sea Dreaming, Small World Music Inc.) was presented. This served as a control condition, in which both participants perceived the same stimuli and were instructed not to interact with each other.

##### Cooperation and competition

The cooperative and competitive computer games were adapted versions of the tasks of Cui et al. (*5*), modified for use in children (*7, 31*). Each player manipulated a cartoon figure of a dolphin by pressing a computer key to either catch a ball together (cooperation) or catch the ball faster than the other player (competition). The cooperative and competitive games were both organized into two task blocks of 20 trials each, interspersed with 30 s rest blocks: rest1, task1, rest2, task2, rest3.

Each trial started with the appearance of the two dolphins on the screen. After 2 s, a black circle appeared in the middle of the screen above the dolphins, the ‘ready’ signal. After a variable interval of 0.6 - 1.5 s, the circle was replaced by a red-and-white ball, the ‘go’ signal. After the ‘go’ signal had appeared, the dyads were asked to react as simultaneously as possible to win shared points (cooperation) or to react faster than the partner to win points for themselves (competition). At the end of each trial, a feedback screen was shown, followed by a result screen.

In the cooperative task, a dyad won shared points if the difference in the participants’ response times was below T = 1/8 [RT1 + RT2], where RT1 and RT2 represent the response times of the two participants. In successful cooperation trials, both dolphins jumped to the ball (feedback screen, 1.5 s) and caught the ball (result screen, 1.5 s), while in unsuccessful trials only the faster dolphin jumped to the ball (feedback screen) and neither of them caught the ball (result screen). Based on this information, the participants were able to adjust their response times. If one of the players reacted too early, that is, before the ‘go’ signal, the trial was started again from the beginning, and both players lost a point.

In the competition trials, the faster dolphin jumped to the ball (feedback screen, 1 s) and caught the ball (result screen, 1 s). If both participants reacted simultaneously within a margin of error of 50 ms, both jumped to the ball (feedback screen) and caught the ball (result screen). If one of the players reacted too early, again, the trial was restarted from the beginning, and only that player lost a point.

### 4.2 fNIRS preprocessing

First, the raw intensity data were converted into optical density data. Second, motion artifacts were detected using a moving standard deviation and reduced through cubic spline interpolation of the affected part of the signal (*59*). Third, the optical density data were converted into HbO and HbR concentration changes using the modified Beer-Lambert law, in which the differential pathlength factor was individually estimated based on the wavelength and the participant’s age (*60*). Finally, the HbO/HbR time series were detrended using a discrete cosine transform set. Noisy fNIRS channels were identified based on a semiautomated procedure, using several objective criteria (coefficient of variation, ‘flat line’ detection, anticorrelation of HbO/HbR) in combination with visual inspection of the HbO/HbR signals and the wavelet transform plots. Based on this procedure, 7.84% of signal pairs were excluded from further analyses. If more than 25% of a participant’s channels under a specific experimental condition were classified as noisy, the complete fNIRS measurement was excluded (text S4).

### 4.3 Graphs

#### 4.3.1 Connectivity estimator

To formulate the interbrain networks, we first quantified the statistical dependencies between the dyad’s brain signals via the bivariate wavelet coherence [WCO; (*61*)]. The WCO is a widely applied functional connectivity estimator that localizes signal dependencies in the time-frequency space and is appropriate for nonstationary time-series, such as fNIRS signals. For each WCO matrix, the coefficients were aggregated into a scalar representing the connectivity estimator. To increase the robustness of the estimator, we applied padding to reduce broader dispersions and considered only salient WCO coefficients that were higher than a cut-off value, determined on the basis of surrogate time-series, and lay within the cone-of-influence (for more information on the calculation of the WCO, the surrogates and the cut-off value see text S5). Finally, we calculated the percentage of salient values within a task-related frequency band between 0.5 Hz and 0.08 Hz (period length: 2.02 s - 12.80 s) for the baseline condition as well as each of the two cooperative/competitive task blocks. The task-related frequency band was chosen based on previous studies (*7, 31*) and included the trial duration (~7 s for cooperation, ~6 s for competition).

#### 4.3.2 Graph construction

A bipartite graph *G* = (*V*_1_ ∪ *V*_2_, *E*) was constructed from the connectivity between the brain region of participant 1 and the brain regions of participant 2. The nodes comprising the two disjoint sets *V*_1_ and *V*_2_, also termed partite sets, represented the spatially localized sources for participant 1 and participant 2, respectively. *E* ⊆ *V*_1_ × *V*_2_ denotes the edges with the corresponding weights *W*, defined by the connectivity estimator (section 4.3.1). For each dyad and task, a complete bipartite graph *G_complete_* was generated with |*V*_1_| = |*V*_2_| = 22 nodes. Finally, edges from noisy channels were excluded. For further implementation details see text S6.

#### 4.3.3 Graph reduction

To weaken the influence of spurious links and to ensure a more robust network topology, the interbrain networks were reduced using null distributions obtained based on shuffled pairs. A shuffled signal pair was formed from two signals from an adult and child who did not participate in the same experiment. We generated all possible shuffled adult-child pairs and calculated their connectivity. With this blockwise permutation, assuming exchangeability of the participant IDs while holding the channel combination and condition fixed, we created a set of null distributions. Subsequently, the 95% quantile of each null distribution was set as the threshold for the respective channel and condition. Through the consideration of the 95% quantile of each null-distribution, this blockwise permutation of shuffled pairs endowed the choice of thresholds with a more solid theoretical foundation and generated multiple large null distributions for each fixed variable combination. Thus, only edges related to the ‘true’ interaction of the dyad were considered, rather than those related to random or systemic similarities between the brain signals due to the same experimental condition. In the following the reduced graphs are denoted by *G*(*V*_1_ ∪ *V*_2_, *E*^−^), with *G_comptete_* (*V*_1_ ∪ *V*_2_, *E*), *s. t.E*^−^(G) ⊆ *E*(*G_comptete_*).

#### 4.3.4 Graph topology

From the reduced graphs, both global and nodal density indices were derived.

**Global interbrain density,** denoted by *d_global_*, was defined as the number of interbrain links that survived permutation, normalized with respect to the total maximum number of possible links (*62*), after ‘noisy channels’ were excluded.

**Nodal interbrain density,** denoted by *d_nodal_*, was defined as the number of surviving edges for each node of the adult and child participants, again normalized with respect to the total number of possible edges. Thus, the nodal density estimates how strongly the neural signals of a given node are coherent with the signals of all other nodes of the partner and thereby allows the individual contributions of a particular node to the global connectivity to be determined.

#### 4.3.5 Bipartite Erdős-Rényi graphs

To study whether the nodal characteristics differed between the original graphs and graphs with a random nodal structure, we adopted bipartite versions of binomial Erdős-Rényi graphs (*63*). Our adjusted generation procedure creates a rewired undirected graph for each original graph such that the generated binomial bipartite Erdős-Rényi graph has the same global density as the interbrain network and only the nodal properties are disturbed.

In this rewiring process, we define the density-corrected bipartite Erdős-Rényi graph *G_Reny_*(*n, m, p*) with *n* = |*V*_1_|, *m* = |*V*_2_|, and *p* = *d_global_*(*G*(*V*_1_ ∪ *V*_2_, *E*^−^)). The graph *G_Reny_* (*n, m, p*) is constructed with the binomial bipartite Erdős-Rényi algorithm such that the vertex sets *E*^*^(*G_Reny_*) ⊆ *E*(*G_compiete_*), with *G_complete_* (*V*_1_ ∪ *V_2_*, *E*) have a corresponding interbrain network *G*(*V*_1_ ∪ *V_2_*, *E*^−^). The probability *p* of generating an edge in the set *E*^*^(*G_Reny_*) is set equal to the global density of the corresponding interbrain network *G*(*V*_1_ ∪ *V*_2_, *E*^−^). Hence, this rewiring process preserves the characteristics of the original graph such that *G_Reny_* (*n, m, p*) will have the same global effects as the corresponding interbrain network *G*(*V*_1_ ∪ *V*_2_, *E*^−^).

### 4.4 Non-negative matrix factorization

To jointly learn a low dimensional representation across all graphs, NMF was applied to the vectorized nodal metrics. To this end, we concatenated the nodal densities (*m* = 44 nodes, 22 each for the child and adult) of all graphs (*n=330*) to form the matrix 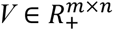. As the nodal density ranges between zero and one, *V* contains only nonnegative entries. This matrix was then approximately factorized into a basis matrix 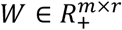 and a coefficient matrix 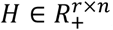, *s. t. V* ≈ *WH* (*17*), where the rank, denoted by *r*, is constrained to (*n* + *m*)*r* < *nm*. The coefficient matrix *H* encodes the nodal densities for each of the *r* components to be used in subsequent analyses. The basis matrix *W* provides the assignment of nodes to components, thus, can be understood as a ‘dictionary’ in which to look up which nodes contribute how strongly to a component. Due to the non-negative constraint, the individual components are strictly additive, making them naturally interpretable, with higher values indicating higher INS. Furthermore, each node can contribute to several components with different weights.

To obtain stable results, we performed 10000 iterations of each NMF procedure. The aim of NMF is to minimize the reconstruction error, which generally describes the differences between the original matrix and the product of the decompositions *W* and *H*. In this study, the reconstruction error was calculated based on the Frobenius norm because it plausibly assumes the noise of the reconstruction to be Gaussian and permits efficient optimization with a coordinate descent solver (*64*). Moreover, the NMF algorithm was initialized via nonnegative double singular value decomposition to ensure a sparse and meaningful representation (*65*).

### 4.5 Inference for interbrain networks

#### 4.5.1 Bayesian hierarchical model

To analyze task and partner effects on global density (section 2.1), a BHM was calculated with the global density as the dependent variable and competition (0 = baseline, 1 = competition), cooperation (0 = baseline, 1 = cooperation) and partner (0 = stranger, 1 = mother) as well as the two-way interactions between task (competition/cooperation) and partner as predictors. For the analysis of nodal effects (section 2.3), an equivalent multivariate BHM was fitted, which included all four NMF components as response variables. The unconditional main effects of competition, cooperation and partner are reported across partner and task conditions, respectively.

#### 4.5.2 Model implementations and quality checks

The BHM analyses were formulated and checked in accordance with the following procedure. Prior to the analyses, missing values were imputed via multiple imputation (text S8). After visual inspection of the data and their distributions, models and priors were specified. We chose default priors that were non- or weakly informative, thus having negligible influence on the results [for more information on the priors, see (*66*)]. Second, the models were fitted with three chains, each of which contained 5000 samples, with samples 1 to 2000 serving as warmup samples to calibrate the sampler, thus yielding a total of 9000 posterior samples. The joint posterior parameter distributions were approximated using the No U-Turn Sampler [NUTS; (*67*)], a type of Markov chain Monte Carlo sampler. Third, to ensure an unbiased inference process, the convergence of the models was checked using the 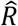 and effective sample size (ESS) indices as well as by visually inspecting the trace plots for regions in the parameter space that could not be efficiently explored by the sampler (funnels). All chains had converged 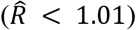, and the ESS values were sufficiently large to accurately derive the quantile intervals of the posterior distribution (table S1). Fourth, posterior predictive checks were performed by plotting the observed data alongside the predicted data sampled from the posterior predictive distribution to examine how well the model reflected the actually observed data. The predictive accuracies of the models were further assessed using leave-one-out cross-validation [LOO; (*68, 69*)] and the Watanabe-Akaike information criterion [WAIC; (*70*)]. In all models, lognormal response distributions were chosen since these yielded better fits than Gaussian distributions. A value of 0.1 was added to the nodal densities prior to the analyses because the logarithm of zero is not defined. First, maximal models were fitted, with intercepts and experimental effects allowed to vary by child (called ‘group-level effects’). Afterwards, group-level effects were reduced backwards in a stepwise fashion, and models of different complexity were compared via the LOO criterion. Maximal models showed the best fit and are therefore reported in the paper. Of note is that the here reported results showed no meaningful differences between the maximal model and the reduced model, in which only the intercept varied by child.

In the results section, for each effect of interest, we reported the mean of its estimated marginal posterior distribution as well as its two-sided 90% CI, also termed the ‘highest posterior density interval’, which is defined as a probabilistic interval that is believed to contain a given parameter (*71*). Further implementation details can be found in text S8.

### 4.6 Classification of interbrain networks

#### 4.6.1 Embeddings and classifiers

State-of-the-art embeddings based on four different methodological approaches were implemented. All embeddings were trained in an unsupervised fashion on graphs constructed as described in section 4.3.

##### A. Skipgram based embeddings

Skipgram refers to a feedforward neural network architecture that was originally applied in natural language processing. Graph2Vec (*27*) is a skipgram-based embedding that decomposes each graph into nonlinear substructures by extracting a subgraph for each node via a Weisfeiler-Lehman kernel. These subgraphs are then used in a skipgram model to learn a lower-rank representation in which graphs composed of similar root subgraphs have similar vector representations. Additionally, we applied ‘Graph and Line graph to vector’ [GL2Vec; (*28*)], which uses the edge-adjacency matrix to reformulate the underlying graphs as line graphs and allows the incorporation of additional attributes of nodes as features. We set the bipartite color, i.e., which nodes belong to the child and which belong to the partner, as a feature.

##### B. Spectral based embeddings

Spectral based embeddings attempt to generalize convolutional operations to non-Euclidean graph domains. Here, we applied the geometric scattering embedding (*29*), which performs (signal-) processing on graphs (*25*) via the wavelet scattering transform. Accordingly, diffusion wavelets calculated from the adjacency matrix and degree matrix are applied to the node signals. These node signals encode a series of node attributes that are captured during random walks across the graph. Again, we adopted the bipartite color as a node attribute.

##### C. Structural role-based embedding

The local degree profile [LDP; (*30*)] is a simple embedding that is calculated based on five topological features: the nodal degree and the mean, minimum, maximum and standard deviation of the nodal degree across a 1-neighborhood. These nodal features are mapped into *n* = 32 bins of uniform width. These bins encode each graph into an *n*-dimensional vector.

##### D. Our network representation

As described in section 4.4, we generated a matrix, representing the nodal densities per node and dyad, which was then decomposed via NMF to obtain a low-rank representation of the reduced bipartite graphs. The resulting coefficient matrix was log transformed, due to its heavy-tailed distribution.

Due to graph reduction (see 4.3.3), deconfounded graphs may contain isolated nodes, i.e., nodes without any edges. However, most kernel-based and neural-network-based embeddings require a connected graph to ensure that each node of the graph can be reached (weak connectivity assumption). Hence, embeddings A and B were computed based on the connected weighted bipartite graphs (see 4.3.1) but with adjusted connectivity estimators to ensure a fair comparison. The connectivity estimators were block centered with respect to the mean of the shuffled pairs in the corresponding experimental condition, task block and channel combination (as described in section 4.3.3) to mitigate the bias induced by noninteractional synchrony. For further details, see text S9.

The embeddings were then input into classifiers. Here, we applied a L2-regularized logistic regression classifier (ridge classification) (*35*), Gaussian process classifier (*36*), and a support vector machine with a linear kernel (*37*). We chose these classifiers since they are well established and represent different linear classification approaches.

#### 4.6.2 Hyperparameter optimization

Hyperparameters indirectly adjust the parametrization of elements of the machine learning algorithms, such as the regularization term of a cost function. Hence, the choice of hyperparameters can strongly influence the outcome of machine learning algorithms (*72*). Here, we optimized the hyperparameters separately for each instance of the cross-validation procedure (section 4.6.3) using Bayesian optimization (*73*). In the case of a moderate parameter space (e.g., *R^d^* with *d* ≤ 20), Gaussian-process-based Bayesian optimization may converge faster to a lower bound solution than alternative approaches (*74*). The hyperparameters were optimized based on the AUC-ROC. Further details can be found in text S9. A detailed parameter space for each classifier and embedding can be found in table S3.

#### 4.6.3 Performance evaluation via repeated, stratified, nested k-fold cross-validation

To ensure a generalizable quantification of the predictive classification performance, we applied k-fold cross-validation to the HbO-based graphs as a resampling strategy. In this implementation, the feature matrix *F* was randomly split into *k* = 5 mutually exclusive subsets of *F* of approximately equal size, termed folds [for the choice of *k* see also (*75*)]. The folds denoted by *F*_1_,*F*_2_,…,*F_k_*, were stratified to preserve the class proportions within each fold. For each fold, preprocessing was applied (see table S3). The hyperparameter optimization was performed separately in a nested loop via stratified cross-validation (*k_nested_* = 3) within each fold of the outer cross-validation. This nested scheme could avoid leakage effects of the training and test sets caused by the optimization process (*76*), which we empirically confirmed (fig. S3). For each instantiation *t* ∈ {1,2,…, *k*}, the classifier was trained and optimized on the disjoined set *F\F_t_* of the fold *F_t_* and the feature matrix *F* and tested on *F_t_*. For each *F_t_*, three performance, namely, ROC-AUC, precision and recall (also termed sensitivity), were calculated (see Fig. 4 and fig. S4). The cross-validation procedure was repeated ten times (*n* = 10) with shuffling to overcome potential bias from the random-partitioning for k-fold cross-validation. The means (*μ*) and standard deviations (*σ*) of the performance metrics for each repetition and fold were compared.

#### 4.6.4 Out-of-sample performance evaluation via the hold-out set

In contrast to the cross-validation procedure, in which the classifiers were trained based on subsets of graphs constructed from the HbO signals, we also trained classifiers based on all graphs constructed from the HbO data (*n* = 330). Next, these trained classifiers were used for prediction based on the graphs constructed from the HbR signals (*n* = 330). This hold-out set was strongly isolated from all previous training procedures. The postprocessing of the embeddings was analogous to that applied in the cross-validation procedure. The classification performance was measured using the same metrics as in the cross-validation procedure. It should be noted that while this out-of-sample prediction procedure benefits from more training and testing samples compared to the cross-validation procedure, the classifiers also suffer from the challenges arising from prediction across datasets. Importantly, HbO and HbR signals have inverse characteristics, and the magnitude of HbR signals is lower, making this classification task challenging.

## Acknowledgements

This project was financially supported by the Excellence Initiative of the German federal state and governments (ERS Seed Fund, OPSF449; granted to K.K, C.B. & D.B.) and the START-Program of the medical faculty of the RWTH Aachen University (granted to V.R.). K.K. received additional funding from the Deutsche Forschungsgemeinschaft (DFG, German Research Foundation) – Project-ID 431549029 – SFB 1451. The Hitachi NIRS system was supported by a funding of the DFG (INST 948/18-1 FUGG). Further, C.G. acknowledges the support by the Helmholtz School for Data Science in Life, Earth and Energy and the computing resources granted by the RWTH Aachen University under project number “0411”.

## Author contributions

C.G., V.R., and K.K. conceived the research project. C.G. and V.R. analyzed and interpreted the data. C.G. drafted the manuscript and V.R. supervised the drafting. All authors revised the manuscript. All authors acquired funding for the research project.

